# Genomic Analysis of Diverse Environmental *Acinetobacter* Isolates Identifies Plasmids, Antibiotic Resistance Genes, and Capsular Polysaccharides Shared with Clinical Strains

**DOI:** 10.1101/2023.10.18.562937

**Authors:** Liam Tobin, Veronica M. Jarocki, Johanna Kenyon, Barbara Drigo, Erica Donner, Steven P. Djordjevic, Mehrad Hamidian

## Abstract

*Acinetobacter baumannii*, an important pathogen known for its widespread antibiotic resistance, has been the focus of extensive research within its genus, primarily involving clinical isolates. Consequently, data on environmental *A. baumannii* and other *Acinetobacter* species remain limited. Here, we utilised Illumina and Nanopore sequencing to analyse the genomes of ten *Acinetobacter* isolates representing six different species sourced from aquatic environments in South Australia. All ten isolates were phylogenetically distinct compared to clinical and other non-clinical *Acinetobacter* strains, often tens of thousands of SNPs from their nearest neighbours. Despite the genetic divergence, we identified p*dif* modules (sections of mobilised DNA) carrying clinically important antimicrobial resistance genes in species other than *A. baumannii*, including carbapenemase *oxa58,* tetracycline resistance gene *tet*(*39*) and macrolide resistance genes *msr(E)-mph(E).* All of these p*dif* modules were located on plasmids with high sequence homology to those circulating in globally distributed *A. baumannii* ST1 and ST2 clones. The environmental *A. baumannii* isolate characterised here (SAAb472; ST350) did not possess any native plasmids; however, it could capture two clinically important plasmids (pRAY and pACICU2) with high transfer frequencies. Furthermore, *A. baumannii* SAAb472 possessed virulence genes and a capsular polysaccharide type analogous to clinical strains. Our findings highlight the potential for environmental *Acinetobacter* species to acquire and disseminate clinically important antimicrobial resistance genes, underscoring the need for further research into the ecology and evolution of this important genus.

**IMPORTANCE:** Antimicrobial resistance (AMR) is a global threat to human, animal, and environmental health. Studying AMR in environmental bacteria is crucial to understand the emergence and dissemination of resistance genes and pathogens, and to identify potential reservoirs and transmission routes. This study provides novel insights into the genomic diversity and AMR potential of environmental *Acinetobacter* species. By comparing the genomes of aquatic *Acinetobacter* isolates with clinical and non-clinical strains, we revealed that they are highly divergent yet carry p*dif* modules that encode resistance to antibiotics commonly used in clinical settings. We also demonstrated that an environmental *A. baumannii* isolate can acquire clinically relevant plasmids and carries virulence factors similar to those of hospital-associated strains. These findings suggest that environmental *Acinetobacter* species may serve as reservoirs and vectors of clinically important genes. Consequently, further research is warranted to comprehensively understand the ecology and evolution of this genus.

## 1. INTRODUCTION

*Acinetobacter* is a genus of Gram-negative bacteria comprising over 85 species with different ecological niches and clinical impacts. Some *Acinetobacter* species are benign and ubiquitous in nature (e.g., found in soil, water, and animals), while others are pathogenic and frequently found in clinical settings (1–5). *Acinetobacter baumannii* is the most prevalent and problematic species in healthcare settings (6, 7), causing a range of nosocomial infections, including pneumonia, as well as bloodstream, urinary tract and wound infections (8). A major challenge in treating *A. baumannii* infections arises from its remarkable ability to acquire resistance to multiple antibiotics, facilitated primarily by horizontal transfer of antibiotic resistance genes via mobile genetic elements (MGEs), such as transposons and plasmids (7, 9–12). Due to its clinical relevance and high levels of antimicrobial resistance, *A. baumannii* has been the subject of extensive research in the last two decades (8, 13).

Pathogenic *Acinetobacter* species have several virulence genes that aid in evading host immune system responses and increase survival and spread throughout its host. For example, the *ompA* gene encodes a key antigenic factor (OmpA) that enables immune evasion (14, 15). Other factors, such as iron acquisition systems and serum/complement resistance, also facilitate *in vivo* survival (14, 15). Moreover, *Acinteobacter* spp. produce a variety of complex carbohydrate structures on their cell surface, such as capsular polysaccharide (CPS), lipooligosaccharide (LOS), and/or lipopolysaccharide (LPS) with an O-antigen covalently attached to the outer core (OC) moiety of the LOS (16). These structures are important virulence factors for many Gram-negative bacteria. In *Acinetobacter*, surface polysaccharides have been studied most extensively in *A. baumannii*, where it has been established that the species produces CPS and LOS (16). The genes that direct the synthesis of CPS and the OC component of the LOS are diverse and can vary even between closely related strains belonging to the same sequence type (ST). Typing these genes is considered an important primary step in strain characterisation (11, 16, 17).

The environment is an important yet understudied reservoir of resistant bacteria, despite the knowledge that prominent extended-spectrum β-lactamases (ESBLs), quinolone resistance genes and carbapenemases originated from marine and soil bacterium and subsequently entered clinical isolates through plasmids (18–20). Given the wide environmental spread of *Acinetobacter* strains, there is a growing consensus that One Health issues – which recognise that humans, animals, and the environment are interconnected – must be addressed through a comprehensive, integrated research approach (21, 22). This involves studying strains of *Acinetobacter* that have been isolated in clinical settings alongside those found in the natural environment to better understand the complex relationships and genetic exchange events between and within each niche (23). However, to date, the primary focus of comparative genomics research has been on hospital-acquired *A. baumannii* strains. As a result, the evolution, genetic structure, virulence determinants, antimicrobial resistance genes, and their associated MGEs present in environmental strains, particularly those not belonging to the *baumannii* species, are poorly understood and remain largely unexplored.

In this study, we performed genomic analyses on ten *Acinetobacter* isolates recovered from South Australian aquatic samples. We show that, while the isolates were genetically unrelated to clinical strains, they share common antibiotic resistance p*dif* modules. This study provides new evidence that environmental strains might act as reservoirs for some of the clinically significant antibiotic resistance genes.

## 2. MATERIALS AND METHODS

### 2.1 Sample collection and isolation of *Acinetobacter* species

Influent wastewater and lake samples were collected as described previously (24). Briefly, water samples (∼10 L) were collected in triplicate, stored on ice and processed within 2 – 3 hours post collection. All samples were plated, in triplicate, on Oxoid Brilliance CRE agar plates [ThermoFisher Scientific, Australia] after 10-fold serial dilutions, using 500 µL from 2 – 3 consecutive dilutions. All cultures were incubated at 25, 37, and 44°C for 24 hours. Single colonies growing on CRE agar were picked up and streaked on plates counting agar (PCA) [ThermoFisher Scientific, Australia]. PCA cultures were then incubated at 37°C for 18 – 24 hours. *Acinetobacter* isolates were initially identified by matrix-assisted laser desorption ionisation-time of flight mass spectrometry (MALDI-TOF MS) using Bruker Daltonics, operated in linear positive mode under MALDI Biotyper 3.0 real-time classification v3.1. Isolates were stored in glycerol (40% v/v) at –80°C.

### 2.2 Antibiotic resistance profile

The resistance patterns of 23 antibiotics, including meropenem (10 µg), imipenem (10 µg), ampicillin (25 µg), cefotaxime (30 µg), ceftriaxone (30 µg), ceftazidime (30 µg), ampicillin/sulbactam (10/10 µg), tobramycin (10 µg), gentamicin (10 µg), spectinomycin (25 µg), netilmicin (30 µg), kanamycin (30 µg), amikacin (30 µg), neomycin (30 µg), streptomycin (25 µg), sulfonamide (100 µg), rifampicin (30 µg), trimethoprim (5 µg), nalidixic acid (30 µg), ciprofloxacin (5 µg), florfenicol (30 µg), chloramphenicol (30 µg), and tetracycline (30 µg) were determined using the standard disc diffusion and standard microbroth dilution methods, which were previously described (**25**). The resistance patterns were analysed based on the guidelines of CLSI for *Acinetobacter* spp. and the calibrated dichotomous sensitivity (CDS) disc diffusion test (http://cdstest.net) in cases where a CLSI breakpoint was not available. This was applicable for antibiotics such as netilmicin, streptomycin, spectinomycin, sulfamethoxazole, nalidixic acid and rifampicin.

### 2.3 Plasmid transfer assays

Transfer studies were performed using a rifampicin mutant of *A. baumannii* SAAb472 (named SAAb472^rif^), which was made in this study as previously described, as a recipient. *A. baumannii* ACICU, which carries the conjugative plasmid pACICU2 (GenBank accession number CP031382) was used as donor in conjugation assays (26). Conjugation assays were conducted using the traditional mating assay at 37°C on agar upon an overnight culture of donor and recipient cells as previously described (27). Briefly, equal amounts of overnight cultures of the donor (ACICU) and recipient (SAAb472^rif^) were mixed and incubated on an L-agar plate overnight. Cells were re-suspended and diluted in 0.9% saline, and transconjugants were selected by plating on MHA plates containing rifampicin (100 mg/L) and kanamycin (100 mg/L). Transfer frequency (transconjugants/donor) was the average of 3 determinations. Potential transconjugants were purified and checked for growth on L-agar containing kanamycin (20 mg/L) and tobramycin (10 mg/L), to which the donor was resistant and the recipient susceptible. Transformation assays were done using electroporation as we previously described (28) and plasmid DNA purified from *A. baumannii* D36 (GenBank accession number CP012952), which carries the small plasmid pRAY* (GenBank accession number CP012954). Transformation frequency was calculated as transformants per µg plasmid DNA.

### 2.4 Whole genome sequencing (WGS), genome assembly, quality control (QC) and annotation

Short read sequencing was performed using the Illumina Nextseq500 platform and reads were assembled using Shovill v1.1 (https://github.com/tseemann/shovill) with –-trim option and the default SPAdes assembler (29). Resulting draft genomes were QCed using assembly-stats v1.0.1 (github.com/sanger-pathogens/assembly-stats). Long read sequencing was performed using the Nanopore GridION platform and consensus long-read assemblies were achieved using Trycycler v0.5.4 (30) in conjunction with Flye v2.9.2 (31), miniasm/minipolish v0.1.3 (32), and Raven v1.8.1 (33) assemblers. To create accurate hybrid assemblies, consensus long read assemblies were polished with short-reads using Polypolish v0.5.0 (34) followed by POLCA (35). All genomes were annotated using the Prokka pipeline v1.14.6 (36) with the –-compliant and –-addgenes options.

### 2.5 Sequence analysis and screening for antibiotic resistance and virulence genes

Resistance genes and insertion sequences were annotated manually using ResFinder (https://cge.cbs.dtu.dk/services/ResFinder/) and ISFinder (https://isfinder.biotoul.fr), respectively. Sequence types were determined using the Institut Pasteur Multi-Locus Sequence Typing (MLST) scheme using the mlst v2.22.1 (https://github.com/tseemann/mlst). The sequence of a set of genes known to be associated with virulence in *A. baumannii* (37) was used to screen the genomes. *Kaptive v. 2.0.5* (38) was initially used to detect genes for the surface polysaccharides, capsular polysaccharide (CPS; K) and the outer-core (OC) component of the lipooligosaccharide (LOS). Command-line searches utilised the available *A. baumannii* K locus (KL) (39) and OC locus (OCL) (40) reference sequence databases, which include 241 KL and 22 OCL, respectively. The minimum identity cut-off parameter for tBLASTn gene searches conducted by *Kaptive* was set to 60%. Protein coding regions were characterised using BLASTp (blast.ncbi.nlm.nih.gov/Blast.cgi?PAGE=Proteins) and Pfam (pfam.xfam.org) searches. Standalone BLAST was used to further characterise the structure of plasmids. The SnapGene® v6.0.5 software was used to manually annotate regions of interest and draw figures to scale using the Illustrator® v26.2.1 program. EasyFig *v. 2.2.5* (41) was used to generate KL and OCL sequence comparisons.

### 2.6 Phylogenetic analysis

Core genome alignments (Block Mapping and Gathering with Entropy and recombination filtered) were produced using Panaroo v1.3.2 (42) with –-clean-mode strict and –a core options, and fed into IQ-Tree2 v2.2.0.3 (43) to produce Maximum Likelihood phylogenetic trees using –m MFP (best fit determined by ModelFinder) and –B 1000 (bootstrap replicates) options. All trees were visualised using Interactive Tree of Life (iTOL) v5 (44). For comparative analyses, additional genomes (n=351) were downloaded from National Center for Biotechnology Information (NCBI) databases (April 2023) using NCBI Datasets v14.18.0 (github.com/ncbi/datasets). For additional taxonomic classifications, average nucleotide identity BLAST (ANIb) (45) with a > 95% cut-off and *in silico* DNA-DNA hybridisations (DDH) (46) with a > 70% cut-off (Formula 2), were performed for species delimitation using NCBI representative strains as references. Pairwise single nucleotide polymorphisms (SNPs) were determined using snp-dists v0.8.2 (github.com/tseemann/snp-dists) SNP matrix heatmaps were produced in RStudio v4.0.5 using pheatmap and ape packages. Pangenomes were calculated using Panaroo and visualised using Phandango v1.3.0 (47). Genome wide association studies (GWASs) were conducted using scoary v1.6.16 (48) with –-no_pairwise options, in conjunction with a gene presence/absence binary matrix produced byPanaroo.

## 3. RESULTS

### 3.1 Phylogenetic analysis reveals significant genetic variability among environmental isolates

Ten *Acinetobacter* isolates were recovered from aquatic environments in South Australia, including influent wastewater (IW) and a lake (Table 1). Initially identified using MALDI-TOF MS, the isolates were classified as *A. baumannii* (n = 1), *Acinetobacter towneri* (n = 3), *Acinetobacter johnsonii* (n = 1) and *Acinetobacter* spp. (n = 5). However, while MALDI-TOF MS is widely used for bacterial identification, environmental isolates, especially those from complex ecosystems like wastewater, can exhibit significant genetic variation leading to incorrect speciation or ambiguous results (49). Thus, to confirm MALDI-TOF speciation, a Maximum Likelihood core genome phylogeny was constructed using 80 representative genomes for *Acinetobacter* species available on NCBI (Supplementary Figure 1). Consequently, three of the five isolates that could not be identified beyond the genus level by MALDI-TOF MS, were assigned a species: *Acinetobacter gerneri* (n = 1; ANIb = 97.04%; DDH = 80.20%) and *Acinetobacter chinensis* (n = 2; ANIb = 96.57% and 97.02%; DDH = 74.10% and 76.50%). The remaining two isolates, SAAs470 and SAAs474, formed their own distinct phylogenetic branch and likely represent a novel *Acinetobacter* species.

**Table 1.**
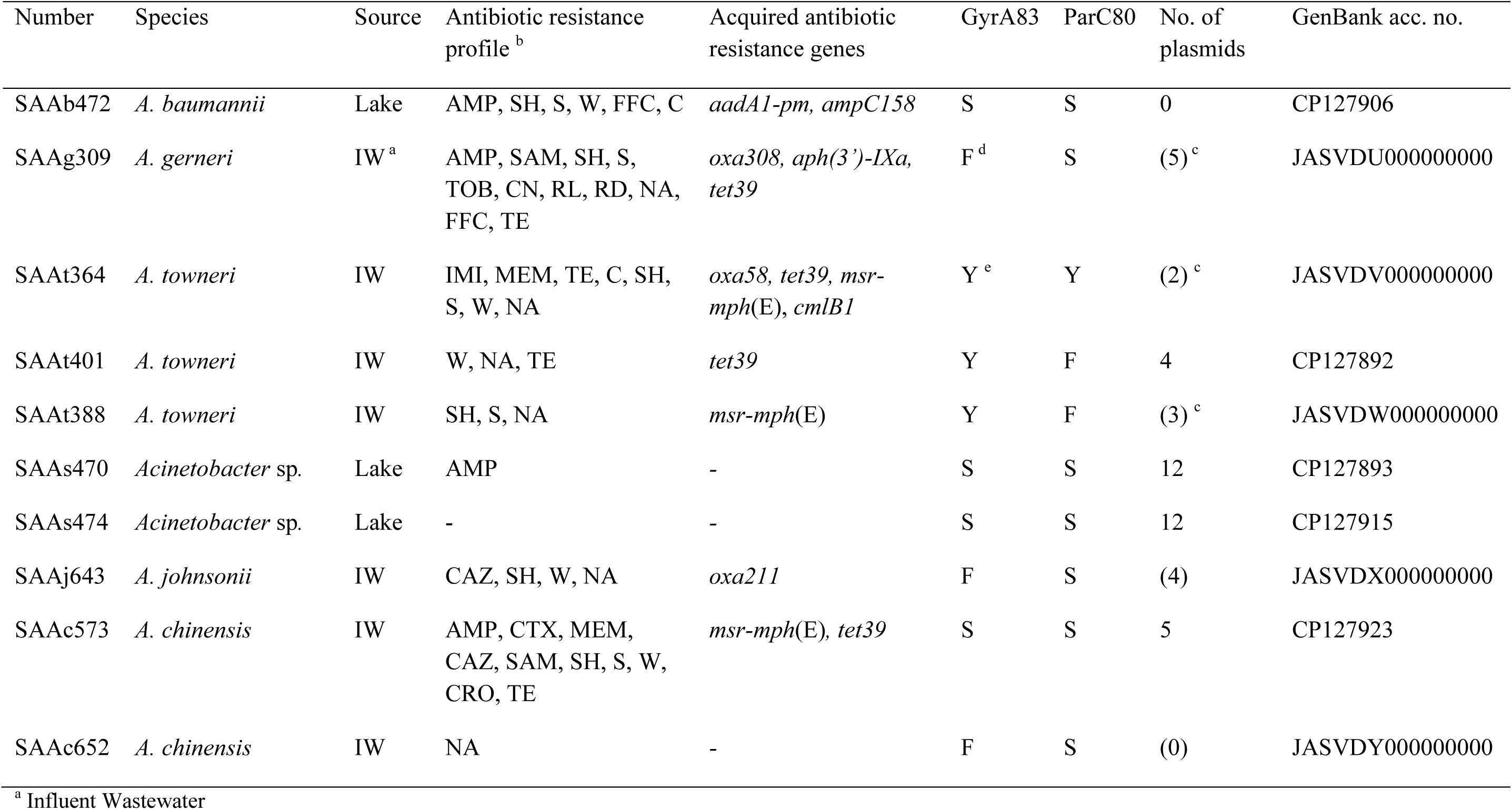
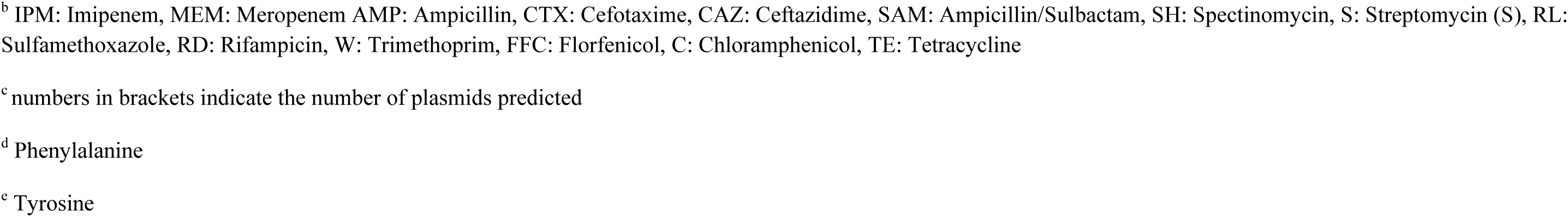
Properties of the *Acinetobacter* strains analysed in this study.

Further phylogenetic trees and SNP heatmaps were built to determine the genetic relatedness of our environmental isolates to *Acinetobacter* genomes previously deposited in public databases. These analyses were not performed for *A. chinensis* and *A. gerneri* due to the limited number of genomes currently available (excluding isolates from this study: *A. chinensis* = 2, *A. gerneri* = 1). For *A. baumannii,* assemblies available for clinical isolates from Australia (n = 128) and global non-clinical isolates (n = 45) were used to construct a Maximum Likelihood core genome phylogenetic tree (Figure 1A). The analysis demonstrated a highly diverse *A. baumannii* population consisting of 26 STs, sharing 2,595 core genes of a total of 10,901 genes (24% of the pangenome), and an average SNP pairwise distance of 23,572 SNPs (Figure 1B). The *A. baumannii* isolate from this study, SAAb472 obtained from a lake, belonged to ST350 and formed its own distinct branch in the phylogenetic tree. SAAb472 was found to be 43,916 SNPs from its nearest neighbour, *A. baumannii* strain GaenseEi-1 (GenBank accession: GCF_002573905) isolated from goose eggshells in Germany 2016.

**FIG 1:**
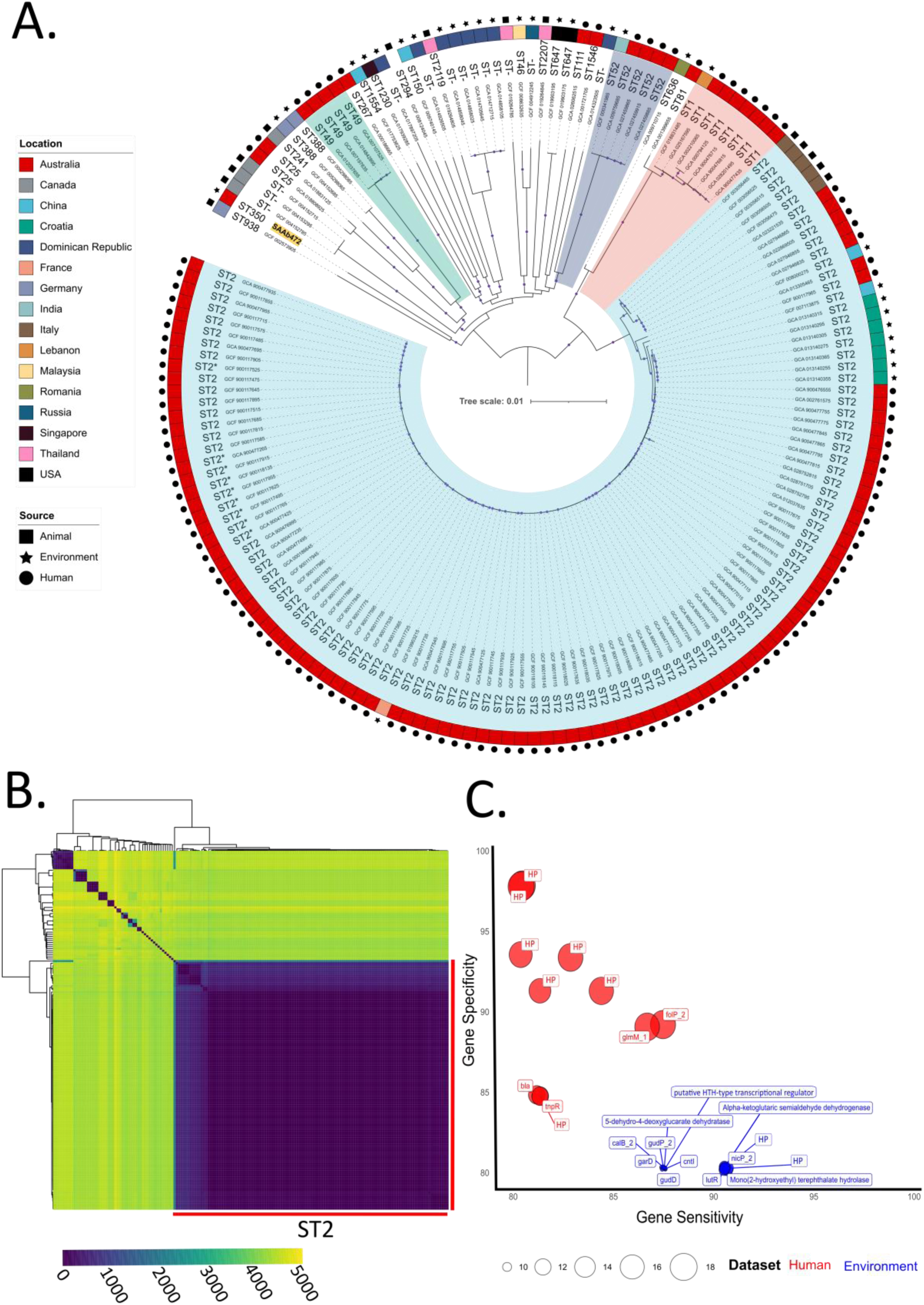
Genetic relatedness of *A. baumannii* clinical, environmental and animal-sourced isolates. A) Maximum Likelihood phylogenetic tree of 174 *A. baumannii* genomes built using a core genome alignment (2,846,563 bp length). The coloured outer ring denotes isolate geographic location and isolate sources are marked as either a circle (human), rectangle (animal) or star (environmental). Tree branches belonging to STs with more than three isolates are highlighted – ST2 isolates in blue, ST1 in red, ST52 in purple, and ST49 in green. Bootstrap values of > 0.95 marked by dots on branches. Isolate from this study is highlighted in yellow. B) Heatmap illustrating pairwise SNP distances between *A. baumannii* isolates. C) Bubble chart depicting highly sensitive and specific genes identified clinical isolates (shown in red) and in environmental isolates (blue). The size of each bubble relates p-values and ranges from 2.31E-10 to 9.58E-19. HP = hypothetical protein.

While the environmental isolates generally displayed higher genetic heterogeneity than human-sourced isolates, even within the clinically significant ST2 population the average pairwise SNPs distance was relatively high at 1170 SNPs (range 0 – 8,525 SNPs). Interestingly, there were two distinct clades of Australian clinical ST2 strains. One clade consisted of 90 isolates with an average distance of 70 SNPs, while the other clade consisted of five isolates that were more closely related to five ST2 isolates from companion animals in Italy than to the larger clade of Australian clinical isolates (2,258 SNPs vs 4,528 SNPs). Also of note was an environmental ST2 strain (GCF_019903215) isolated from a river in France which averaged 120 SNPs from the clonal clade of Australian clinical isolates. These observations are consistent with the ability of *A. baumannii* to spread and adapt to different environments (50). Nevertheless, GWASs did identify genes that were either significantly (p < 0.01) more and less represented in clinical and environmental isolates (Supplementary Data 1), including genes with >80% specificity and >80% sensitivity (Figure 1C). These highly specific and sensitive genes in environmental isolates included those involved in D-glucarate and aldaric acid catabolic processes (5-dehydro-4-deoxyglucarate dehydratase, galactarate dehydratase *garD*, glucarate dehydratase *gudD* and glucarate transporter *gudP*), and mono(2-hydroxyethyl) terephthalate hydrolase, shown to be involved in the degradation and metabolisation of PET plastic for use as a carbon source (51). The genes that were highly sensitive and specific in clinical isolates mostly encoded hypothetical protein (8 of 11 identified), however beta-lactamase *bla*_TEM_ and transposon Tn*3* resolvase *tnpR* were found to be more characteristic of clinical isolates (Figure 1C).

To study the evolutionary context of the *A. johnsonii* isolate, SAAj643 (wastewater), and the three *A. towneri* isolates, SAAt364, SSAt388 and SAAt401 (all wastewater), all assemblies available on NCBI for each respective species were downloaded (*A. johnsonii* = 66; *A. towneri* = 32) and used to build phylogenetic trees (Figure 2A and Figure 2C). *A. johnsonii* SAAj643 formed its own distinct branch and was 50,729 SNPs from its closest neighbour – *A. johnsonii* strain new_MAG-226 (GCA_016790205) isolated from bioreactor sludge in China. On a cautionary note, *A. johnsonii* isolates only shared 294 core genes and 31 (47%) publicly available genomes currently annotated as *A. johnsonii* fall below the ANI threshold for the species. Incorrect taxonomic assignments in the genus *Acinetobacter* are a recognised issue (52), and here we highlight that misclassification may be prominent in *A. johnsonii*, though choice of NCBI representative strains may play a role in low ANIs. Regarding *A. towneri,* there are currently no human-sourced assemblies available on NCBI and most isolates have originated from animal sources (pig and cattle) from China (n = 19; 59%). Like *A. baumannii, A. towneri* isolates were diverse averaging a pairwise distance of 30,445 SNPs (Figure 2D). The wastewater isolates SAAt364, SSAt388 and SAAt401 each formed their own branches in the phylogenetic tree and had an average genetic distance of 27,660 SNPs between them.

**FIG 2:**
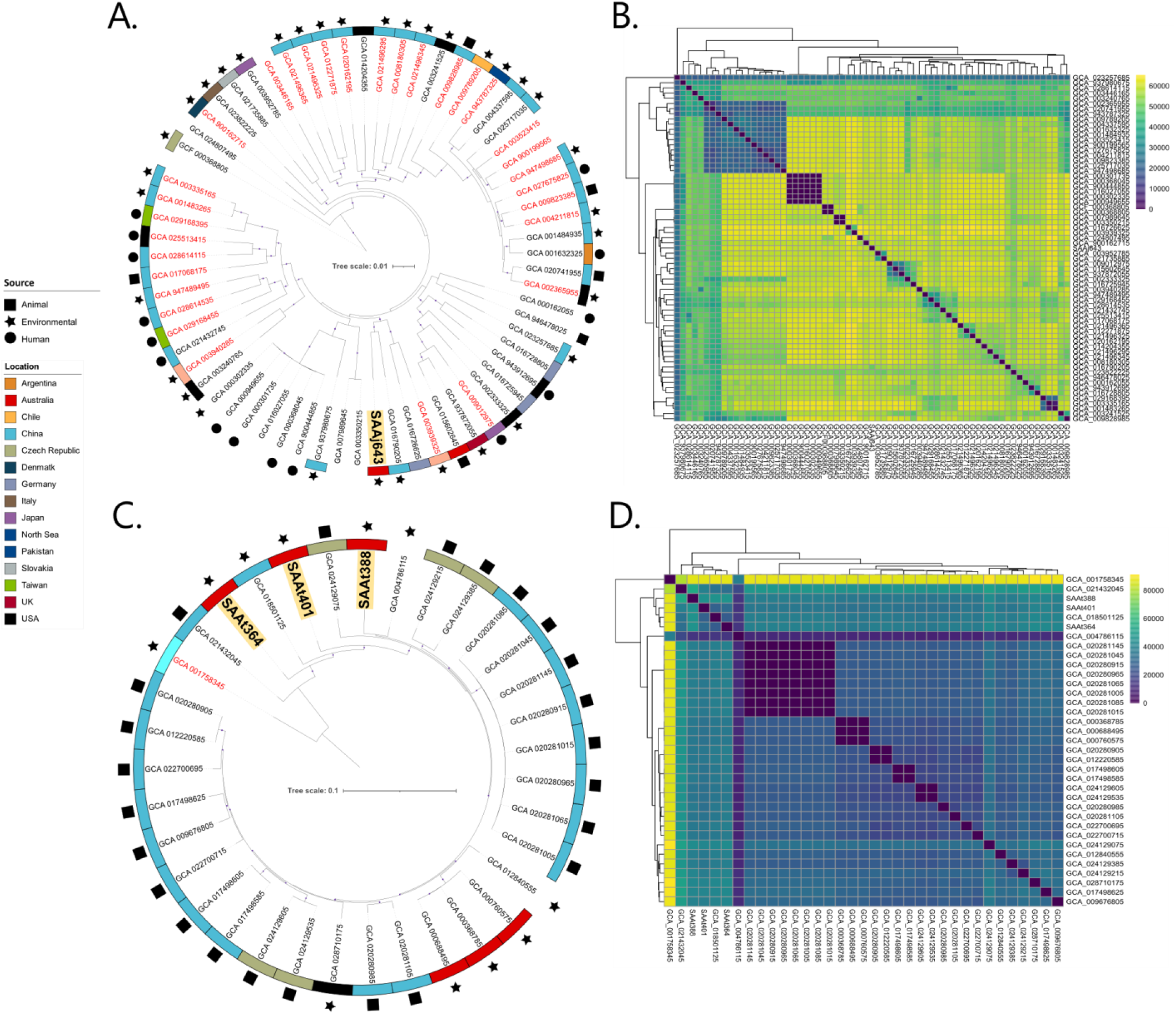
Genetic relatedness of *A. johnsonii* and *A. towneri* isolates. A) Maximum Likelihood phylogenetic tree of 67 *A. johnsonii* genomes built using a core genome alignment (1,948,641 bp length). Isolate from this study in bold and highlighted in yellow. Isolates shown in red mark those that fall under the ANI threshold for an *A. johnsonii* conclusive identification. B) Heatmap illustrating pairwise SNP distances between *A. johnsonii* isolates. C) Maximum Likelihood phylogenetic tree of 35 *A. towneri* genomes built using a core genome alignment (1,758,026 bp length). The three *A. towneri* isolates from this study are in bold and highlighted in yellow. The isolate shown in red fell under the ANI threshold for an *A. towneri* conclusive identification. D: Heatmap illustrating pairwise SNP distances between *A. towneri* isolates.

### 3.2 Antibiotic resistance and genetic context of resistance genes

Except marginal ampicillin resistance, the *Acinetobacter* sp. strains SAAs470 and SAAs474 recovered in a lake, were susceptible to all other antibiotics tested and consistent with this they did not contain any acquired antibiotic resistance genes. However, in all other strains, analysis of the antibiotic resistance profiles indicated resistance to several clinically significant antibiotics, including resistance to carbapenems in *A. towneri* SAAt364 (Table 1), tetracycline resistance in *A. gerneri* SAAg309, *A. towneri* SAAt364, *A. towneri* SAAt401 and *A. chinesis* SAAc573 (Table 1) and aminoglycoside resistance in *A. baumannii* SAAb472 and *A. gerneri* SAAg309 (Table 1). Importantly the *oxa58* carbapenem resistance gene was found in *A. towneri* SAAt364 (Table 1). The *oxa58* gene is one of the most widespread carbapenem resistance genes found in clinical *A. baumannii* strains belonging to the major successful global clones such as ST1, ST2, ST25 etc, (10, 53). This gene is often located on R3-type plasmids surrounded by several complete and partial copies of ISAba2 and ISAba3. Recently, *oxa58* and its companion insertion sequences were shown to be part of a p*dif* module (54). p*dif* modules are frequently found in *Acinetobacter* plasmids. They are made of pairs of Xer recombination sites called plasmid–*dif* (p*dif*) sites that flank a gene or genes (often resistance genes, toxin-antitoxin genes etc) (2, 54). Here, analysis of the draft genome of *A. towneri* SAAt364 showed that *oxa58* is in a p*dif* module like those previously described (54). Analysis of the contig (containing *oxa58*) also suggested that the p*dif* module carrying the *oxa58* carbapenem resistance gene is on an R3-type plasmid closely related to pMAD (with an addition of the *oxa58* p*dif* module) in *A. baumannii* MAD (AY665723). *A. baumannii* MAD is a clinical strain recovered in Toulouse, France, in 2003 (55).

The *tet39* tetracycline resistance gene was found in *A. gerneri* SAAg309, *A. chinensis* SAAc573 and *A. towneri* strains SAAt364 and SAAt401 (see the genetic structure of pSAAt401 in supplementary FIG 4). In addition, the *msr*-*mph*(E) macrolide resistance genes were detected in *A. towneri* strains SAAt364 and SAAt388 and *A. chinensis* SAAc573.

Amongst the genomes completed in this study (n=5), *A. baumannii* SAAb472 did not contain plasmids. However, others, including *A. towneri* SAAt401, *A. chinensis* SAAc573 and the two *Acinetobacter* sp. strains (SAAs470 and SAAs474), carry several diverse novel plasmids, ranging in size from ∼2 to over 150 kb (Table 2).

**Table 2.**
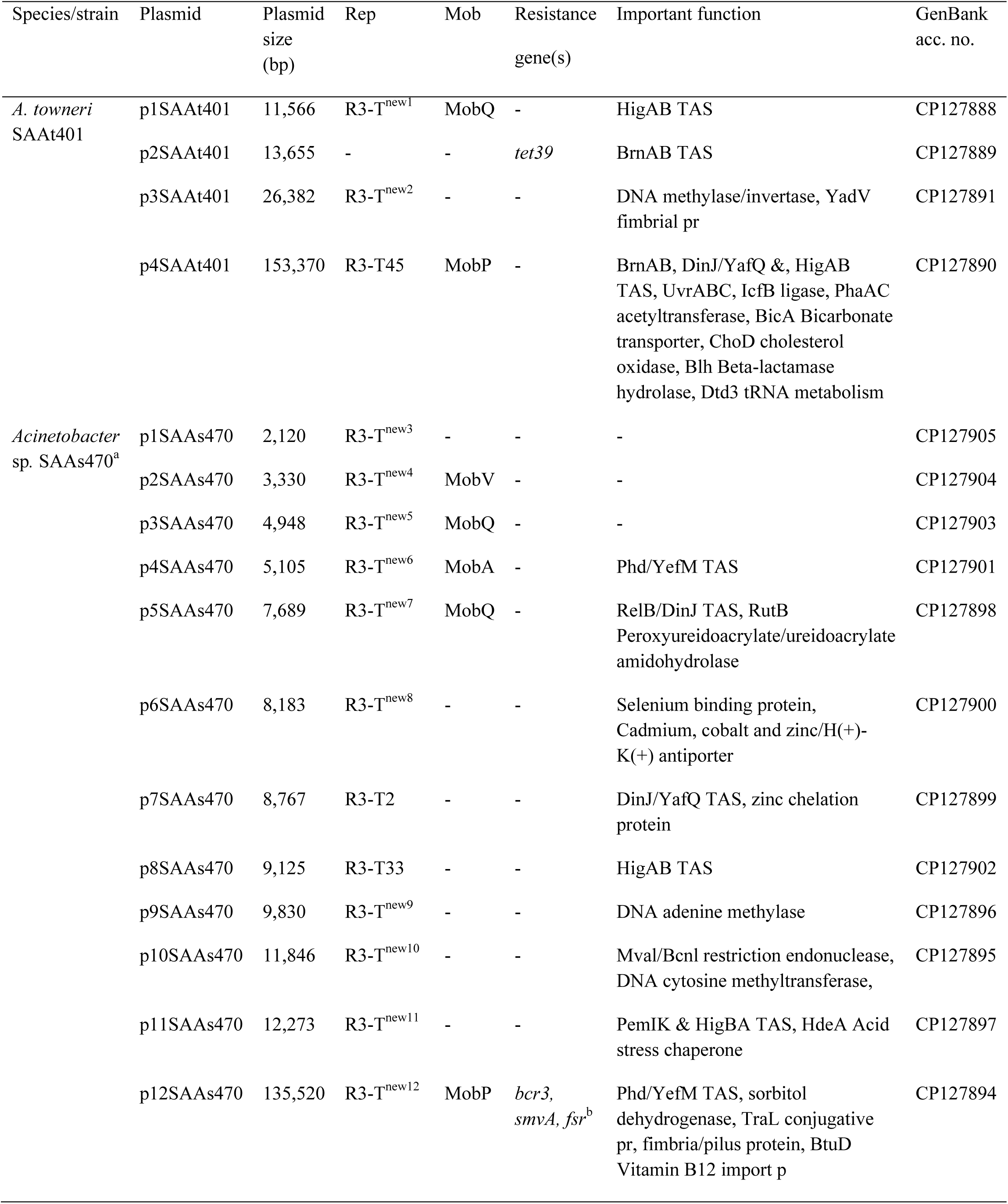
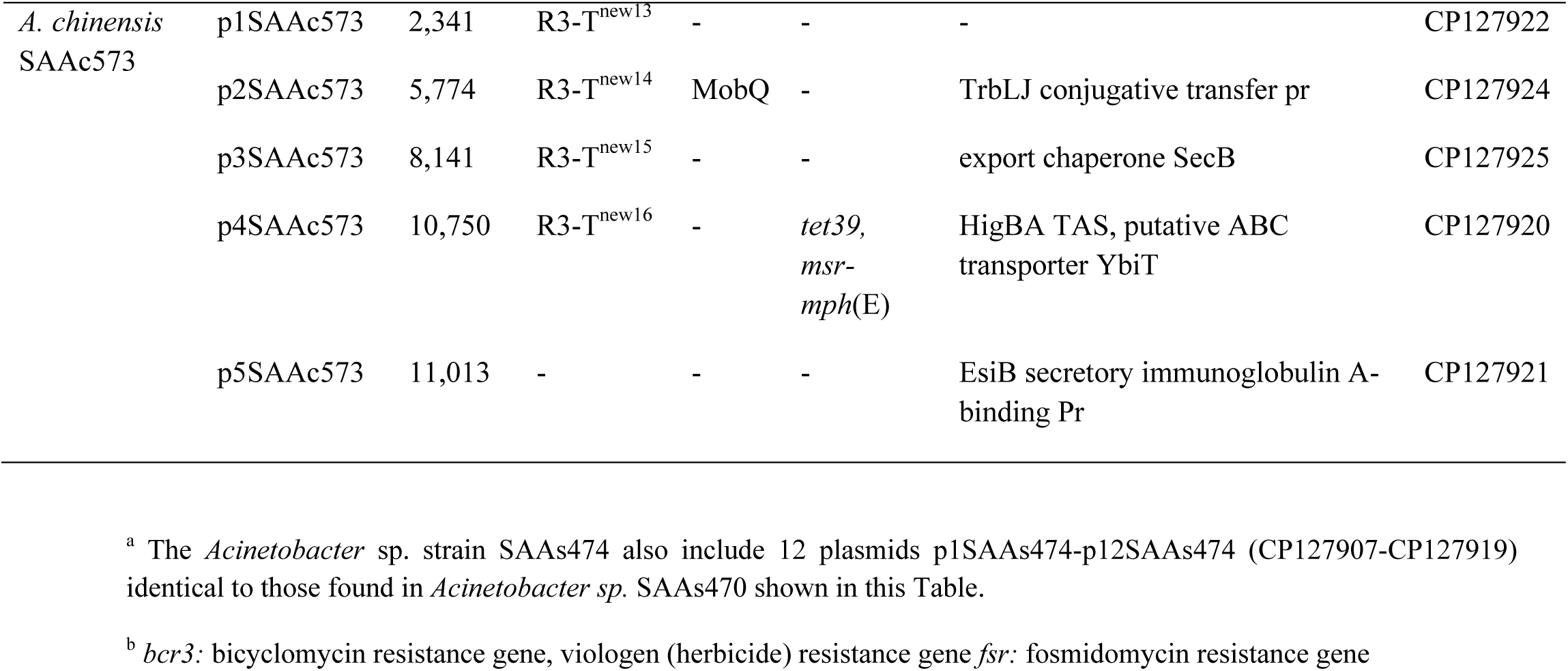
Properties of complete plasmids sequenced in this study.

Notably, all acquired antibiotic resistance genes were found in p*dif* modules located in different plasmid types (Table 2). p*dif* modules are frequently found in *Acinetobacter* plasmids (7, 28, 53, 56). Here, the analysis of the surrounding regions of the *oxa58, tet39* and *msr*-*mph*(E) genes also indicated they are located within p*dif* modules. p*dif* modules containing the *oxa58, tet39* and *msr*-*mph*(E) genes are widespread in strains recovered in clinical samples (53, 54, 57) and in this study we report their presence in a set of environmental isolates indicating that these resistance genes, or the plasmids they are associated with, are able to move between different *Acinetobacter* spp. or onto diverse plasmids.

We also analysed the genetic context of several plasmids that carry the *tet39* and *msr-mph*(E) resistance genes and found several closely related plasmids of both environmental and clinical origins. This indicates that these plasmids can potentially be exchanged between clinical and environmental strains. As an example, analysis of p4AAc573 indicated that it is closely related to multiple plasmids from environmental and clinical strains (Figure 3) particularly p1_010052 (carried by *Acinetobacter* sp. WCHAc010052 recovered in sewage in China in 2015; CP032138), p5637 (carried by *A. baumannii* 2016GDAB1 recovered in a bronchoalveolar lavage in China in 2016; CP065052) and pRCH52-1 (KT346360.1), which was also recovered in a clinical sample in Brisbane, Australia prior to 2012.

**FIG 3.**
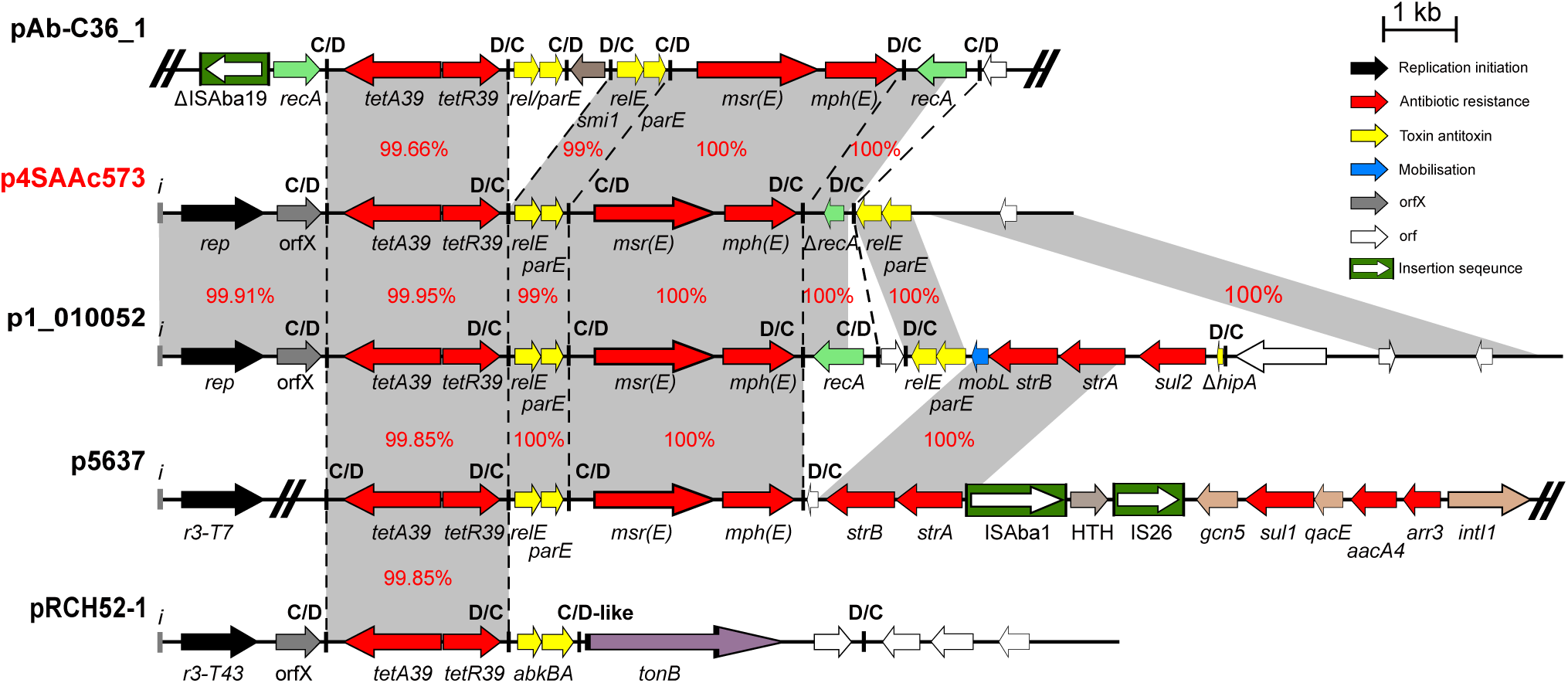
Genetic structure of p2SAAc573 compared to pAb-C36_1, p1_010052, p5637 and pRCH52-1. Filled arrows indicate the orientation and extent of genes. Resistance genes are coloured red, and the filled box coloured green indicate ISAto1. Black arrows are putative replication initiation genes and toxin/anti toxin genes are yellow. Vertical black lines indicate p*dif* sites. Scale bar is shown. SUPPLEMENTARY MATERIAL

In addition, to the closely related plasmids, the *tet39* an *msr*-*mph*(E) p*dif* modules found here were also identified in several unrelated plasmids with clinical origins (FIG 3 and 4) suggesting that apart from plasmids, p*dif* modules can also be readily exchanged between different species and between strains with environmental and clinical origins.

### 3.3 *A. baumannii* SAAb472 can capture clinically important plasmids

*A. baumannii* plasmids play a crucial role in facilitating the spread of clinically significant antibiotic resistance genes. However, there remains a paucity of information concerning the ability of environmental strains to capture these clinically relevant plasmids. Hence, we sought to investigate whether plasmids of clinical importance could be transferred to *A. baumannii* strain SAAb472 (isolated from a lake in South Australia). Two plasmids were used in transfer assays, i) pRAY* (GenBank accession number CP012954.1), a 6 kb plasmid harbouring the *aadB* tobramycin resistance gene (28), and ii) pACICU2 (GenBank accession number CP031382), a 70 kb conjugative plasmid carrying the *aphA6* amikacin resistance gene (58, 59). Both plasmids are widely recognised and prevalent across various *A. baumannii* STs and clinical settings in diverse geographical regions (28, 56, 58, 59). Using electroporation, pRAY was successfully transferred to SAAb472 with a high transfer frequency of 2.99 × 10^7^ CFU/µg. Similarly, pACICU2 was also successfully transferred to SAAb472^rif^ (a rifampicin mutant of SAAb472 generated in this study) through conjugation assays, with a high frequency of 2.1 × 10^-3^.

### 3.4 Environmental strains encode various virulence functions

To understand variations in virulence potential within the South Australian environmental strains, we explored genes that have previously been identified as crucial for virulence in clinical isolates of *A. baumannii*. This included a set of genes encoding various functions such as biofilm formation, outer membrane proteins, capsular surface polysaccharides, regulatory proteins and siderophores (37).

All genes, previously shown to be associated with virulence in *A. baumannii* (2), were identified in the *A. baumannii* isolate signifying that it has the potential to cause infection (Table 3). All genes, encoding regulatory proteins (*bmfRS, gigA* and *gacA/S*) were identified across all genomes studied suggesting the intrinsic nature of these genes within the *Acinetobacter* genus (Table 3). The *ompA* outer membrane gene was absent in *A. towneri* strains SAAt364 and SAAt388 as well as the two *Acinetobacter* sp. strains (SAAs470 and SAAs474) however, the *smpA* and *blc* genes encoding outer membrane proteins were present in all species (Table 3). In contrast, the *bap* gene (associated with biofilm formation) and the *bauA* gene (associated with siderophore production) were absent in all isolates except for *A. baumannii* SAAb472 (Table 3).

**Table 3.**
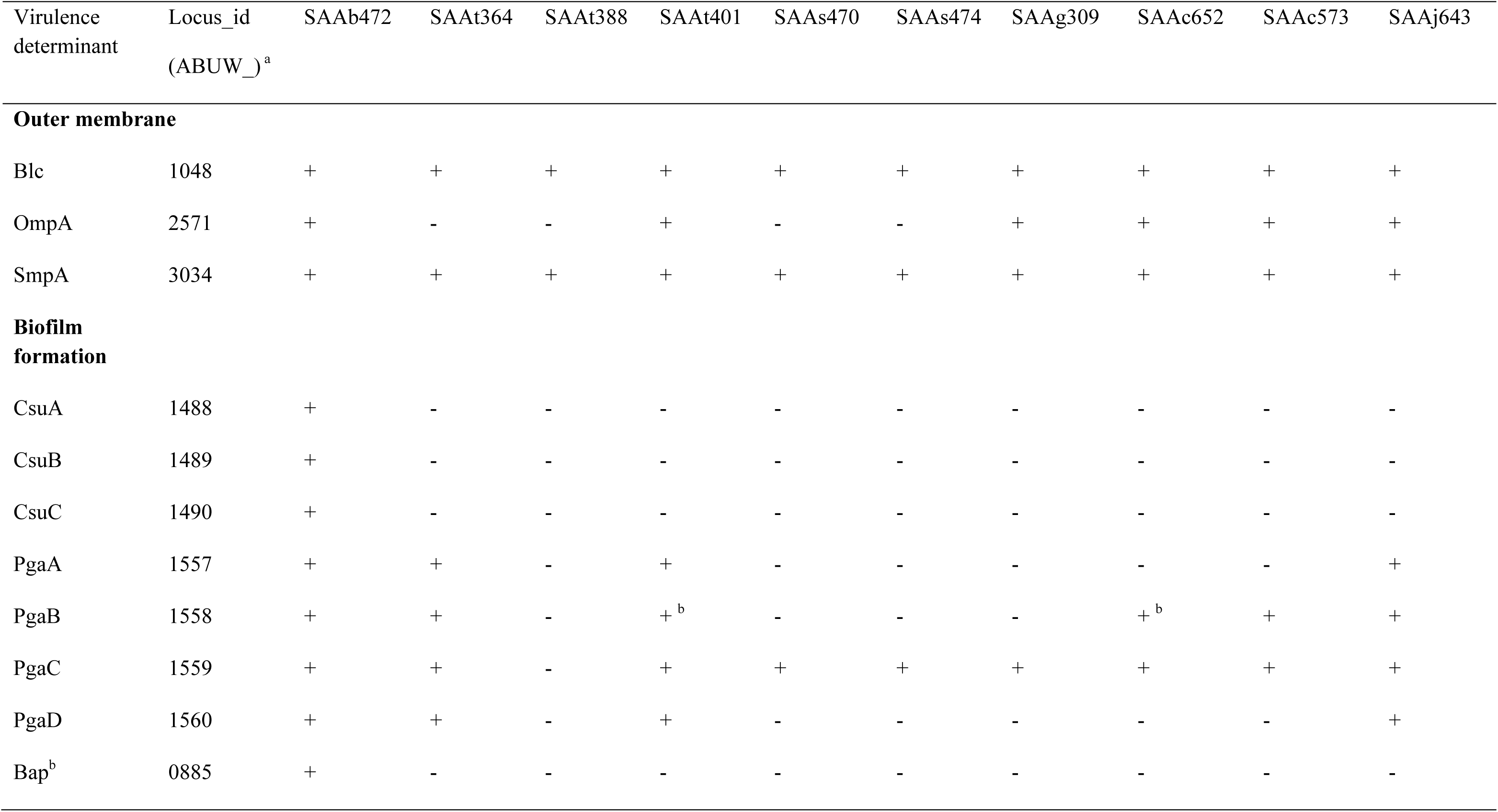
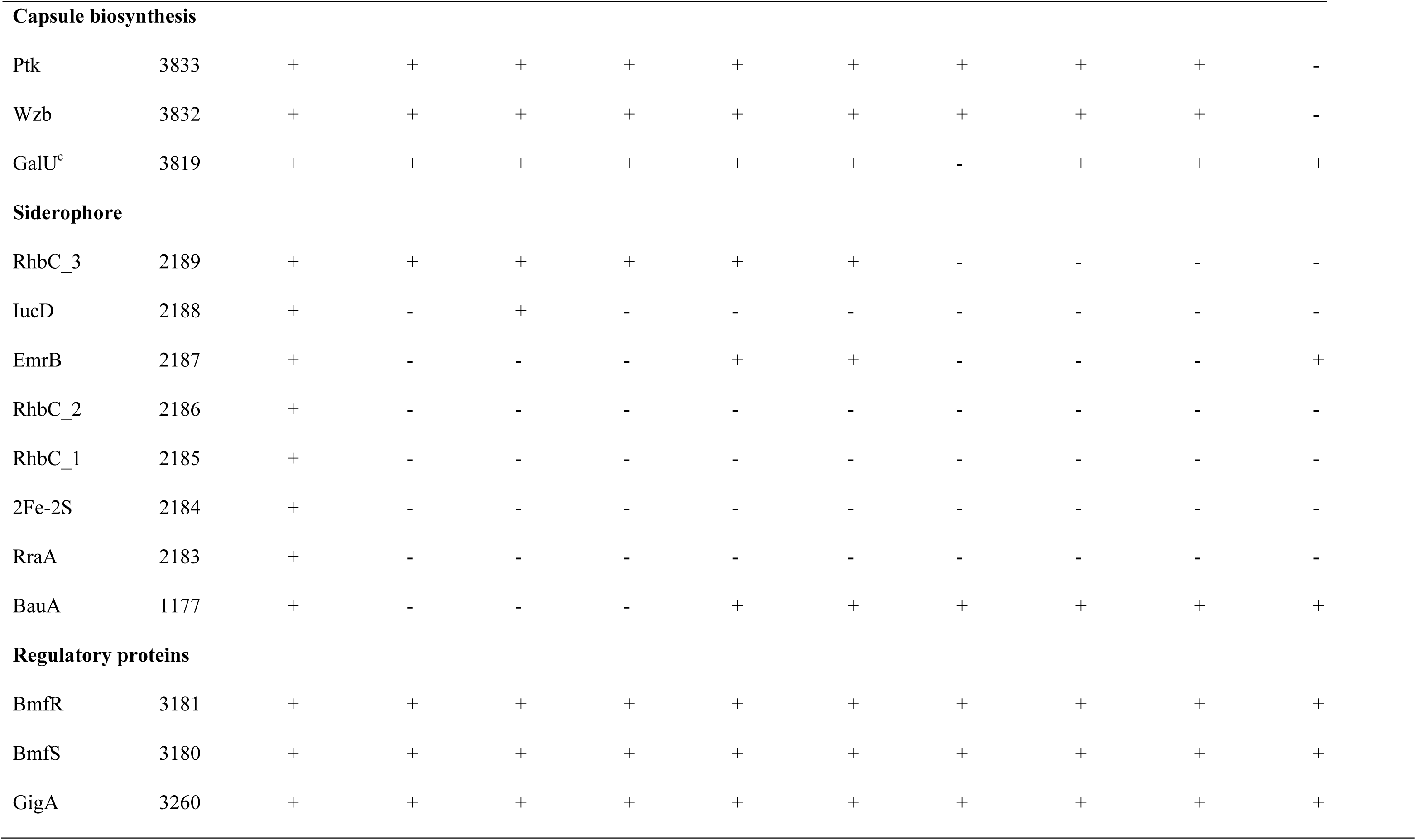
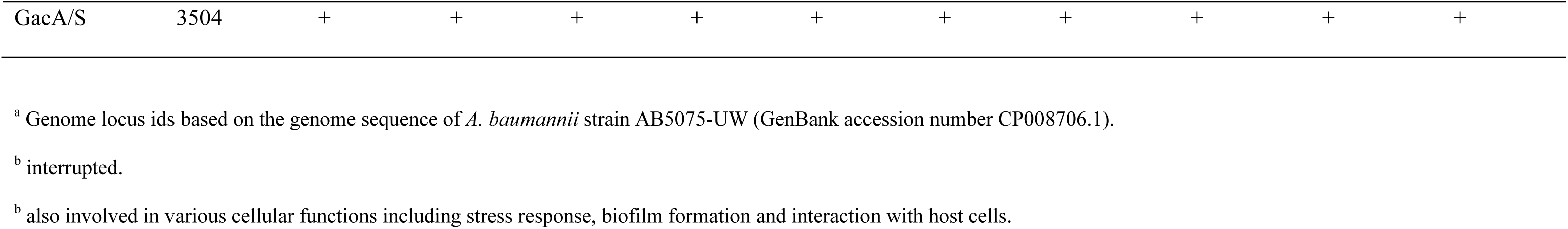
Distribution of virulence functions encoded by South Australian aquatic strains.

Together, while several important virulence determinants were identified across our genomes, there was a high degree of variations even within species. Given that most genomes (9 out of 10) returned hits for common CPS genes, we therefore studied the genetic context and variations within the CPS loci (see below).

### 3.5 Diversity in surface polysaccharide loci

In *A. baumannii*, CPS genes are clustered at the K locus (KL) between conserved *fkpA* and *lldP* genes, whereas genes for the OC of the LOS are at the OC locus (OCL) flanked by *ilvE* and *aspS*. While typing schemes are not currently available for other *Acinetobacter* spp., the K locus of a small number of studied strains have been shown to follow a similar arrangement. Therefore, the ten *Acinetobacter* spp. genomes were screened against the available *A. baumannii* reference sequence databases of 241 KL and 22 OCL to identify common CPS and OC genes to locate the putative positions of polysaccharide loci for further analyses.

Only *A. baumannii* SAAb472 returned a significant match to a KL and OCL reference in the databases. The specific sequence at the K locus was identified as the KL121 CPS biosynthesis gene cluster (100% coverage, 97.93% DNA identity), previously described as carrying *psa* genes for the synthesis of a complex nonulosonic acid known as pseudaminic acid. KL121 has been detected amongst a range of *A. baumannii* isolates from clinical (blood, wound, respiratory) or environmental (white stork, goose eggshells) samples obtained from either USA, Poland, Germany, Brazil or Nigeria. At the OC locus, SAAb472 carries the OCL1 locus type (100% coverage, 98.42% DNA identity) previously found in ∼75% of publicly available *A. baumannii* genome assemblies and common amongst members of the GC1 and GC2 clonal complexes (40).

Significant matches across the entire length of the KL or OCL were not obtained for any of the other *Acinetobacter* genomes. However, a common module of *wza-wzb-wzc* CPS export genes (61-81% translated aa identity) were detected in eight of the nine genomes, identifying the contig(s) and possible location of the K locus. In all eight genomes, this module was found adjacent to other genes encoding products predicted to be involved in polysaccharide biosynthesis. While *wza-wzb-wzc* genes were not found in the SAAj643 genome, other genes associated with CPS synthesis were identified together in a single contig, suggesting that this strain may produce a complex polysaccharide but use different machinery to export it to the cell surface. Each genome was found to harbour a different locus sequence, though the overall genetic organisation of the identified loci (shown in Supplementary FIG 2) resembles that of the *A. baumannii* K locus. Each locus included one or more modules of genes for different sugars, all of which have previously been identified in *A. baumannii* KL.

The putative OC locus in the non-*baumannii* genomes was identified via the finding of genes related to OCL genes from *A. baumannii*. A novel locus sequence was present in all genomes, though the two *Acinetobacter* spp. carried the same locus type (Supplementary FIG 3). As for *A. baumannii*, all loci identified consisted of 1-2 modules of genes for sugar biosynthesis, as well as acetyl-/acyltransferase gene(s) and/or several glycosyltransferase genes. Most included *rmlA, rmlB, rmlC* and *rmlD* genes for synthesis of L-rhamnose or *rmlA/rmlB* for a related sugar. However, like the K loci, very little sequence similarity was observed between the strains.

## 4. DISCUSSION

Environmental bacteria are ubiquitous and play important roles in biogeochemical cycles and ecological interactions. However, they are often intrinsically resistant to a range of antibiotics and constant selective pressures facilitate the acquisition of additional antibiotic resistance genes through horizontal gene transfer. Thus, studying environmental bacteria in the context of AMR is crucial to understand the mechanisms, dynamics and drivers of AMR emergence and spread, as well as to identify potential sources and reservoirs of AMR genes. Here, through a genomic analysis of ten environmental *Acinetobacter* isolates from South Australia we found that, despite being phylogenetically very distinct, several *Acinetobacter* species shared AMR associated MGEs previously only associated with clinical *A. baumannii* strains.

Previous studies have highlighted the high genetic diversity of *Acinetobacter* populations (60, 61). However, the presence of antibiotic resistance genes in the environmental isolates underscores the significance of disseminating resistance determinants in the environment. It is known that the presence of residual antibiotics and other pharmaceutical substances stimulate the bacterial SOS system and promote hypermutation and horizontal gene transfer (HGT) (62). These substances are released into rivers and streams during manufacture and when antibiotics are metabolised. These residues also accumulate in municipal wastewater and urban runoff creating selection pressures in the environment that foster antibiotic-resistance. These processes, in turn, increase the risk of antibiotic resistance acquisition and evolution (63). Additionally, the coexistence of diverse antibiotic resistance genes within a diverse population of pathogens and environmental bacteria, along with elevated rates of HGT, creates a conducive environment where new resistance gene arrangements or mutations can emerge due to these selection pressures. The identification of diverse plasmids harbouring antibiotic resistance genes, including those associated with carbapenem resistance, suggests the mobility and adaptability of resistance elements across various strains. The presence of similar plasmids in both clinical and environmental strains indicates the potential for cross-species transfer, which has implications for public health interventions and strategies to combat the spread of antibiotic resistance. p*dif* sites are remarkable as they are associated with the movement of many genes, including antibiotic and heavy metal resistance genes. p*dif* modules carrying carbapenem, tetracycline and macrolide resistance genes (e.g., *tet39*, *mph*-*msr*(E), *oxa58* and *oxa24*) are widespread in major global clones of *A. baumannii* (28, 54). Here, the identification of the *tet39*, *mph-msr*(E) and *oxa58* p*dif* modules that are often associated with plasmids circulating in globally distributed *A. baumannii* clones (e.g., ST1, ST2, ST25 etc.) is of particular significance. This observation raises concerns about the potential exchange of resistance genes between environmental and clinical strains, further emphasizing the need for holistic One Health approaches in studying the evolution and transmission of antibiotic resistance in the genus.

The successful transfer of clinically significant plasmids to the environmental *A. baumannii* strain SAAb472 highlights the adaptability of environmental strains to acquire plasmids of clinical importance. This combined with the identification of virulence determinants within the environmental isolates, particularly the *A. baumannii* strain SAAb472, underscores the potential of the environmental strains to cause infections in clinical settings.

The diversity in surface polysaccharide loci observed in this study reflects the complexity of virulence factor expression within *Acinetobacter* species. While some genes were shared among strains, the high degree of variation within species emphasizes the need for further investigation to understand the functional implications of these variations in terms of host interactions and environmental adaptations. The fact that CPS loci in other species are arranged in the same general genetic organisation as sequences at the *A. baumannii* K locus may assist with recombination of the central region that determines the structural type driving CPS diversity. However, the KL and most OCL identified in this study were found at different locations in the genome in other species.

In this study, environmental Australian *Acinetobacter* strains were examined, providing valuable insights into genetic diversity, antibiotic resistance, plasmid dynamics, and virulence determinants. The findings not only shed light on the potential role of these strains as reservoirs for antibiotic resistance genes but also emphasise the interconnectedness of environmental and clinical strains. The identification of clinically significant genes within environmental strains underscores the potential risk of gene transfer between different niches, emphasizing the importance of ongoing research into the dynamics of antibiotic resistance and virulence factors in *Acinetobacter* populations. These insights contribute significantly to our understanding of the complex interactions between bacteria, the environment, and human health. They also provide a solid foundation for future studies aimed at mitigating the spread of antibiotic resistance.

## Data Availability

The complete genome, plasmid sequences and short read data of all strains reported in this study have been deposited in the GenBank/EMBL/DDBJ database and are publicly available under the BioProject accession number PRJNA949389.

## ACKNOWLEDGEMENTS

Authors declare no conflict of interest.

This research was supported by the Australian Institute for Microbiology and Infection, University of Technology Sydney, Data Generation Grant and the Australian Centre for Genomic Epidemiological Microbiology (Ausgem), a collaborative research partnership between the New South Wales Department of Primary Industries and the University of Technology Sydney.

J.K is supported by an Australian Research Council Future Fellowship (FT230100400). MH is supported by an Australian Research Council DECRA fellowship (DE200100111).

MH conceptualisation. LT, VJ, JK, MH methodology, investigation, formal analysis. BD resources. VJ, MH writing – original draft. VJ, JK, BD, SD, MH, methodology, writing – review & editing.

## Supplemental Material

Supplementary Data 1: Scoary GWAS raw output

Supplementary Figure 1: Maximum-likelihood phylogenetic tree using 80 *Acinetobacter* representative species and ten South Australian environmental isolates.

Supplementary Figure 2: Genetic arrangement of capsular polysaccharide biosynthesis loci

Supplementary Figure 3: Genetic arrangement of loci for synthesis of the outer core of the lipooligosaccharide/lipopolysaccharide.

Supplementary Figure 4: Genetic structure of p2SAAc573 compared to pAb-C36_1, p1_010052, p5637 and pRCH52-1.

## SUPPLEMENTARY MATERIAL

**FIG S1:**
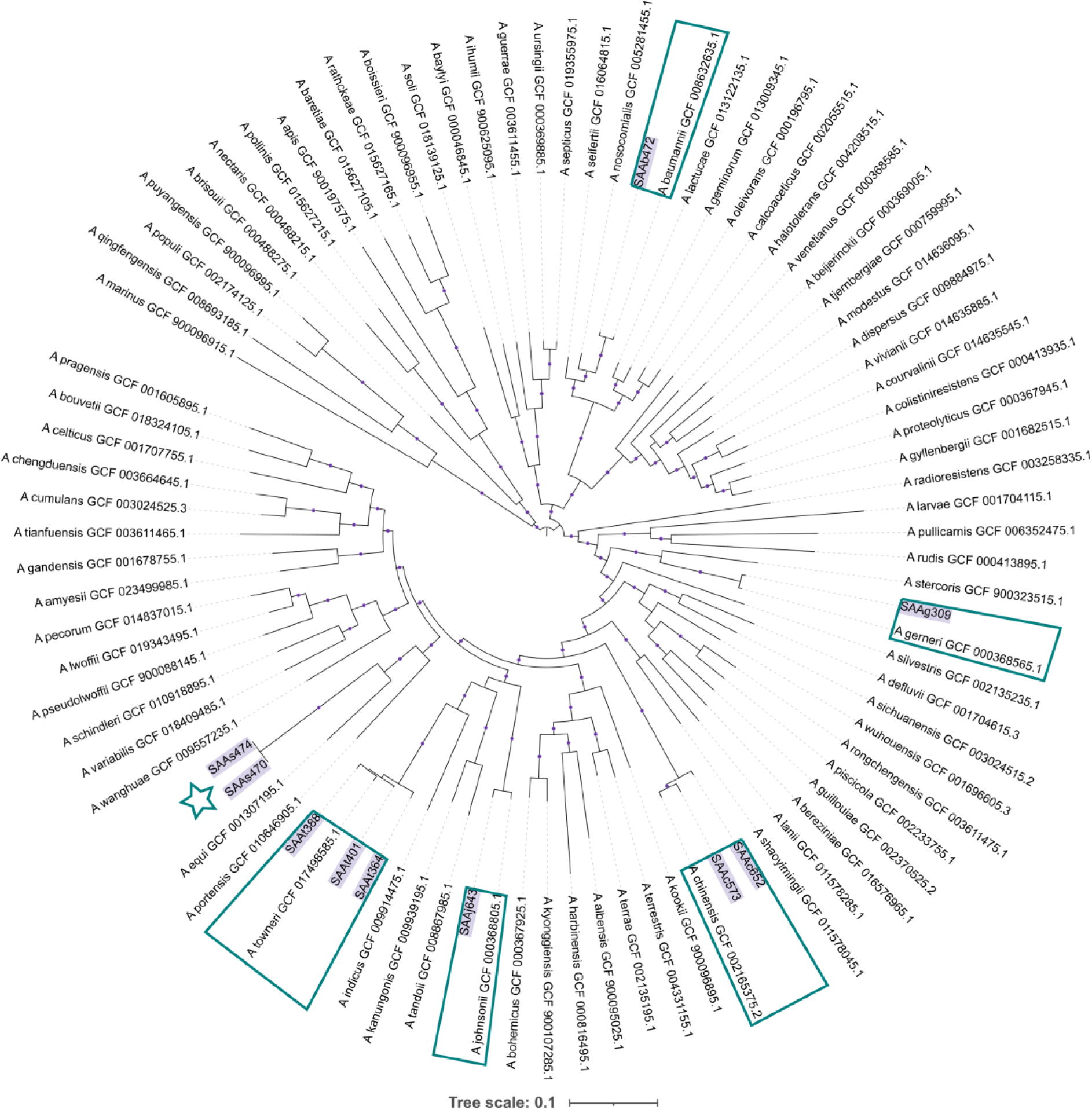
Maximum likelihood phylogenetic tree using 80 *Acinetobacter* representative species genomes and ten South Australian environmental isolates. Tree was built using a core genome alignment (54,841 bp length). Bootstrap values > 0.9 shown as dots on branches. Isolates from this study highlighted in purple and are boxed to include each nearest representative *Acinetobacter* species. The potentially novel species is marked with a star.

**FIG S2:**
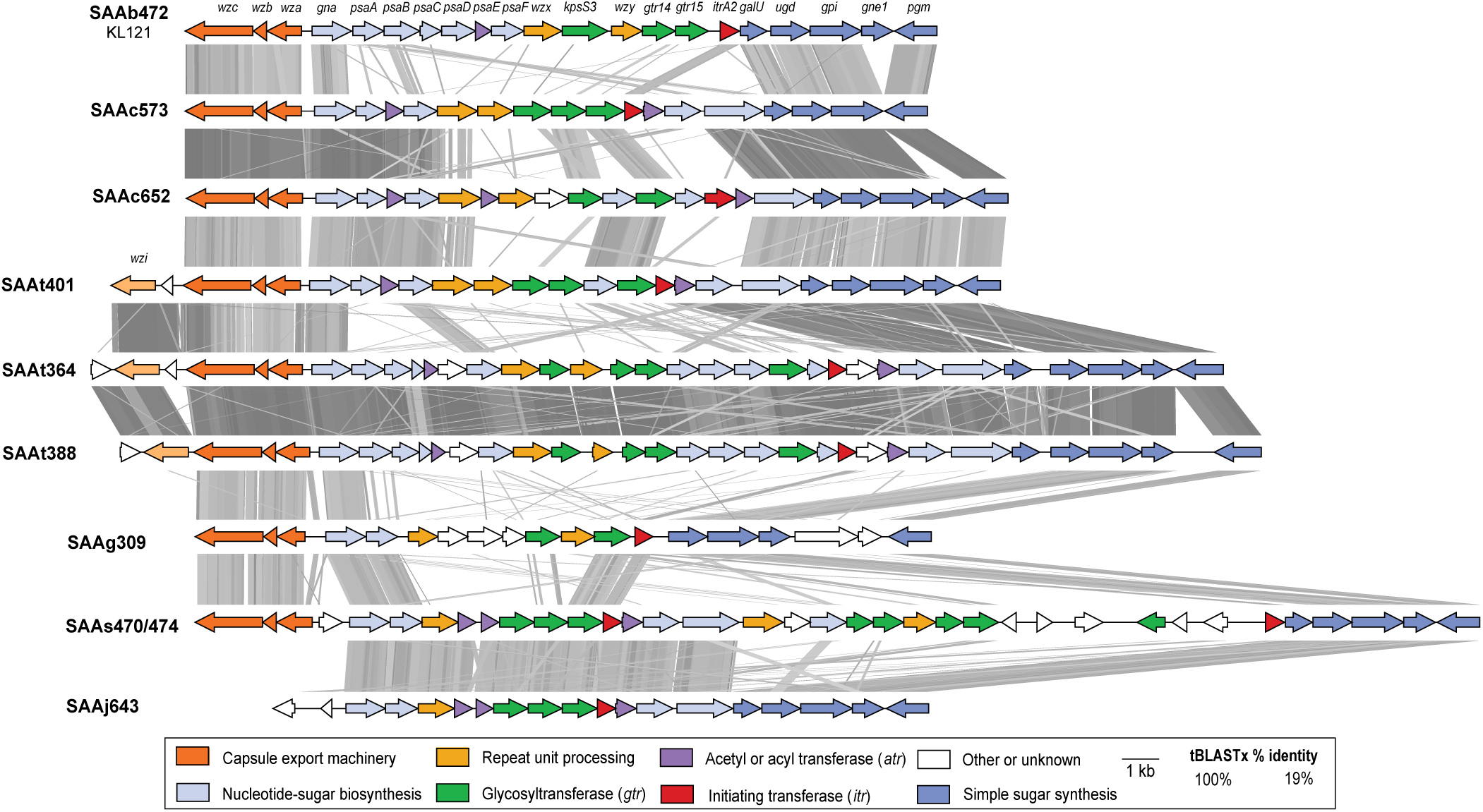
Genetic arrangement of capsular polysaccharide biosynthesis loci identified in the genomes studied here. Genes are shown as arrows indicating the direction of transcription and are coloured by the predicted functional category of their gene products as per legend below. Shading between sequences indicates amino acid sequence identity determined by tBLASTx with scale shown in the legend below. Loci found across more than one contig in SAAg309, SAAt364 and SAAt388 draft genomes.

**FIG S3:**
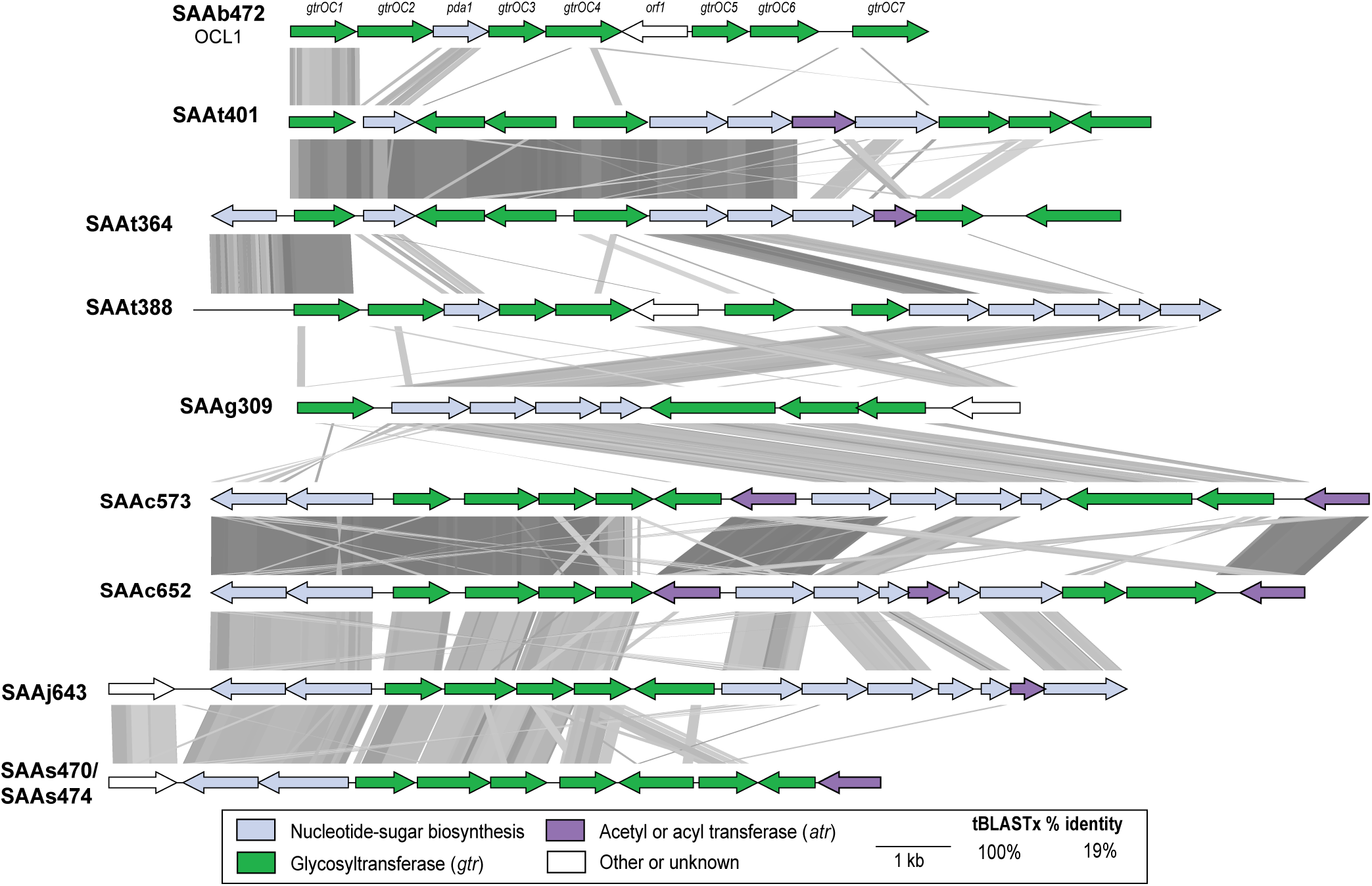
Genetic arrangement of loci for synthesis of the outer core of the lipooligosaccharide/lipopolysaccharide for genomes studied here. Genes are shown as arrows indicating the direction of transcription and are coloured by the predicted functional category of their gene products as per legend below. Shading between sequences indicates amino acid sequence identity determined by tBLASTx with scale shown in the legend below. Loci found across more than one contig in SAAg309, SAAt364, SAAt388 and SAAj643 draft genomes.

**FIG S4.**
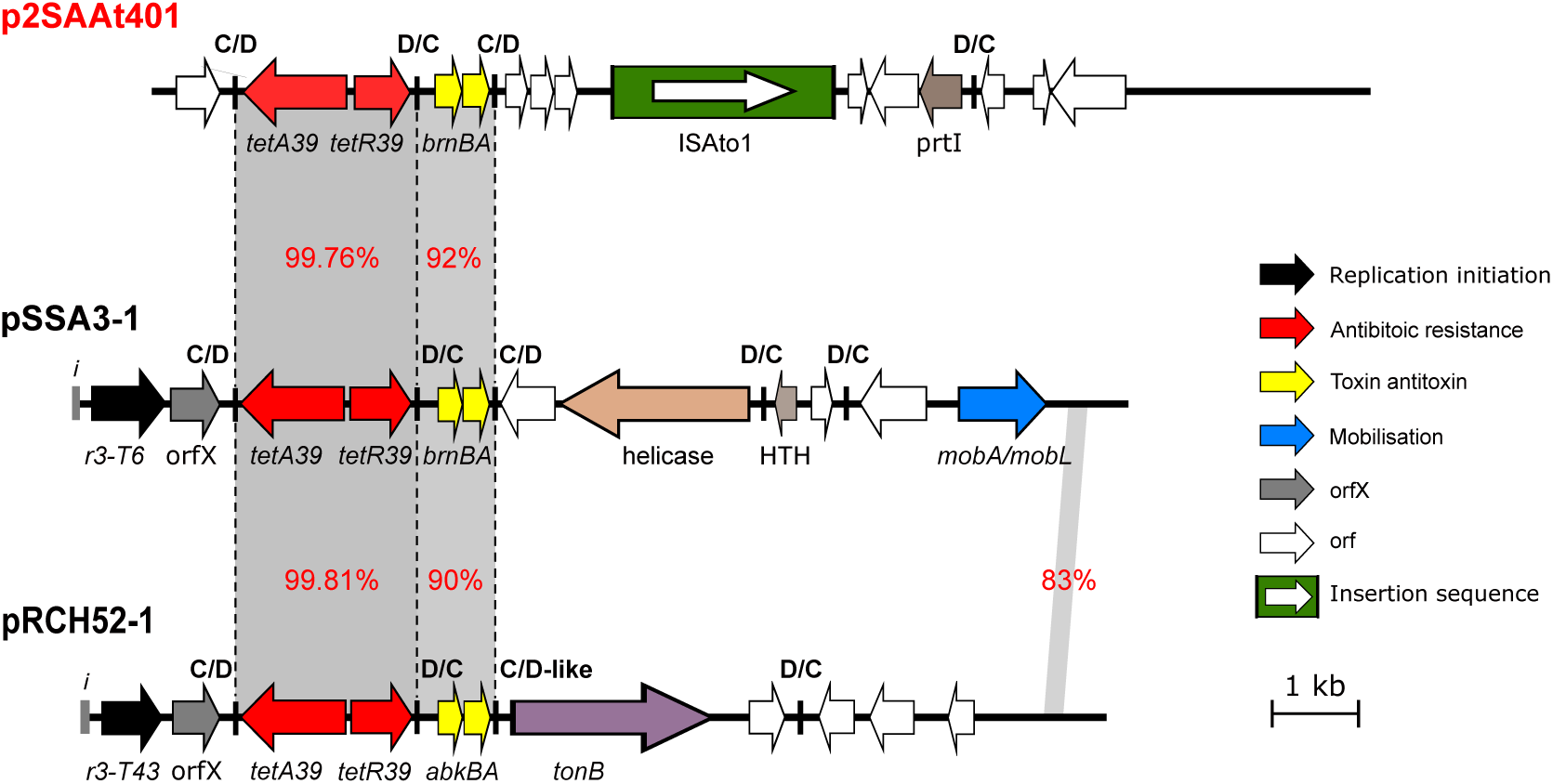
Genetic structure of p2SAAt401 compared to pSSA3-1 and pRCH52-1. Filled arrows indicate the orientation and extent of genes. Resistance genes are coloured red and the filled box coloured green indicate ISAto1. Black arrows are putative replication initiation genes and toxin/anti toxin genes are yellow. Vertical black lines indicate p*dif* sites. Scale bar is shown.

